# Tubule jamming in the developing mouse kidney creates cyclical mechanical stresses in nephron-forming niches

**DOI:** 10.1101/2022.06.03.494718

**Authors:** John M. Viola, Jiageng Liu, Louis S. Prahl, Aria Huang, Trevor J. Chan, Gabriela Hayward-Lara, Catherine M. Porter, Chenjun Shi, Jitao Zhang, Alex J. Hughes

## Abstract

The kidney develops through branching of progressively crowded ureteric bud (UB) tubules at the organ surface. The elongating tubule tips are surrounded by traveling cap mesenchyme niches consisting of nephron progenitors and separated by stromal boundaries. Dynamic interactions between these tissues coordinate a balance between UB tip branching, elongation, and nephron induction that sets nephron numbers for life, impacting the likelihood of adult disease. Such a crowded tissue environment could place geometric limits on the number of niches that can be formed while maintaining mechanical integrity of the tissue. Since space is at a premium, crowding could also force a given niche to prioritize between nephron formation or UB branching differently depending on its spatial context. Here we study the geometric and mechanical consequences of tubule tip crowding at the embryonic kidney surface. Organ curvature reduces and tubule ‘tip domain’ niches pack more closely over developmental time. These together create a semi-crystalline geometry of tips at the kidney surface and a rigidity transition to more solid-like tissue properties at later developmental stages. To infer mechanical dynamics over the branching timescale, we define a new method to infer tip domain ‘ages’ relative to their most recent branch events from fixed kidneys. We find that new tip domains overcome mechanical resistance as they branch and displace close-packed neighbors, transiently increasing mechanical stress in the niche. Ongoing efforts to understand geometric and mechanical effects on niche regulation will clarify variation in kidney tissue composition and advance engineering control strategies for synthetic regenerative tissues.

## Introduction

Tissue-building processes during embryonic kidney development set the number of nephrons and urinary collecting tubules in the adult organ, since no further nephrons are added after postnatal day 4 in mice and around birth in humans ^1^. Starting at ∼E11 in the mouse, the ureteric bud (UB) branches into kidney mesenchyme to form the future urinary collecting ducts. The tubule tips lie at the organ surface during branching and are surrounded by dynamic clusters of cap mesenchyme cells that proliferate and serve as nephron progenitors ^2–6^. Meanwhile signaling pathways including BMP/pSMAD and Wnt/β-catenin induce nephron progenitors to periodically condense and undergo mesenchymal-to-epithelial transition at ‘armpit’ regions beneath UB tips ^7–9^. Nephrons first form as spherical pre-tubular aggregates and later develop into renal vesicles, comma-shaped and S-shaped bodies, simultaneously forming patent lumens with the adjacent UB tip and setting the proximodistal axis of the nephron ^10,11^. While these stages have been characterized anatomically and molecularly ^12,13^, there is less known about temporal coordination between the UB branch cycle, the changing geometry and mechanics in the niche, and nephron condensation events. This is partially explained by temporal and spatial variation in kidney branching morphogenesis. For example, branching of the UB is asynchronous (meaning that UB tips do not branch together at the same time) and asymmetric (meaning that the number of descendants of sister tubules at a given branch generation are different) ^14,15^. Branching asynchrony has effectively led to existing scRNA-seq and imaging data being averaged over tips/niches at different stages in their branch ‘life-cycles’. Therefore, niches associated with ‘young’ tips vs. ‘old’ tips relative to their last branch event have not been separately characterized. Quantitation of niche geometry, mechanics, and signaling over the niche life-cycle are likely necessary to develop a predictive understanding of nephron formation *in vivo* and in engineered systems.

We recently found that mouse embryonic kidney niche geometry is partially predictable through physical modeling of tip repulsion and crowding ^16^. Preliminary analysis of UB tip positions at the surface revealed different crystal-like packing geometries that were at least locally ordered. These observations suggested an analogy to 2D curved crystals of repulsive particles ^17–21^. Such crystals have several fascinating physical properties that could impact the developmental trajectory of the kidney by setting limits on tip and therefore nephron number for a given organ surface area, and by changing the physical microenvironment of tips over their elongation and branching cycles.

Here we combine the theory of curved crystals, rigidity in close-packed systems, and force inference with micromechanical analysis to characterize nephrogenic niche geometry and mechanics. We then peg these properties to the UB branching life-cycle. Our data reveal that niche packing causes a transition from more fluid-to solid-like tissue mechanics that enable mechanical stresses to be sustained among nephrogenic niches after ∼E15. This in turn causes rhythmic cycles of mechanical stress within nephrogenic niches that are synchronized with the UB branching life-cycle. Our work suggests several questions for ongoing study, for example, is nephrogenic signaling activity correlated with rhythmic changes in niche geometry and/or mechanical stress? These data will yield new insight into compositional control of the kidney, which would add to fundamental understanding of disease-relevant variations in nephron endowment ^22–25^, and aid engineering efforts for regenerative medicine.

## Results

### Ureteric bud tubule geometry is partially determined by kidney curvature

During branching morphogenesis, UB tubules duplicate just beneath the kidney surface. Each tubule tip is surrounded by a swarm of ‘cap’ mesenchyme cells that serve as nephron progenitors, with each niche repelling each other ^6,26,27^, creating dense arrays of tip niches separated by thin sheets of stroma (**Fig. 1A, Movie S1**). We previously hypothesized an analogy for close packing of niches in the physics of repulsive or elastic particles at surfaces ^16^. The geometry and mechanics of packing in such systems are affected by particle density and surface curvature ^21,28,29^. We thus began by considering the contribution of kidney curvature to the geometric microenvironment of individual UB epithelial tips. Repulsive particles pack most efficiently on flat surfaces in a hexagonal close-packed (triangular lattice) fashion where each particle has six neighbors (i.e. the coordination number *z* = 6) (**Fig. 1B**). However, wrapping a triangular lattice onto a surface with Gaussian curvature creates an energetic cost that favors the emergence of topological defects called disclinations, in which particles have greater or fewer than six neighbors. Secondly, for curved crystals that are sufficiently large, isolated dislocations are less favorable and are replaced by pairs, clusters, or chains (‘scars’) having ‘excess’ dislocations (**Note S1**) ^17,18,20^. We therefore wondered if UB tip positions adhere to the same topological requirements that set defect number and organization in curved crystals.

**Fig. 1:**
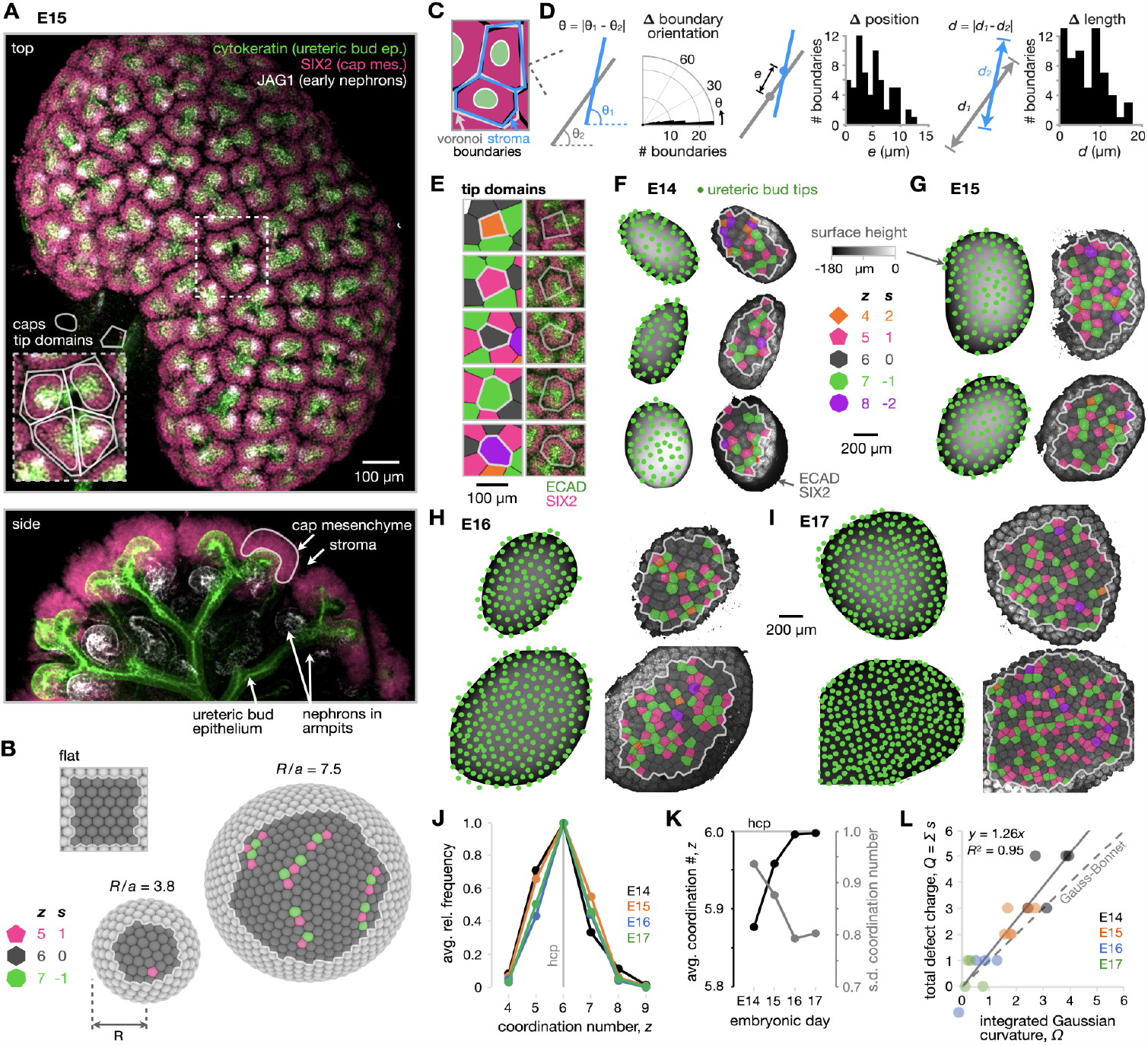
Kidney curvature imposes topological requirements on geometry of close-packed nephrogenic niches at ureteric bud tubule tips. (**A**) Confocal fluorescence maximum projection (top) and xz plane for embryonic day (E)15 mouse kidney. Branching cytokeratin+ UB tubule tips are surrounded by SIX2+ nephron progenitors in the cap mesenchyme. Caps are separated by a thin layer of stroma (unlabeled). JAG1+ early nephrons form at ‘armpits’ beneath tubule tips. We define ‘tip domains’ by Voronoi polygons. (**B**) Schematic of repulsive particle (radius *a*) packing at the surface of a sphere (radius *R*), showing pentagonal and heptagonal topological defects amongst an otherwise hexagonal packing. Defects form pairs, clusters, or chains for *R*/*a* > 5. (**C**) Schematic of overlap between tip domain and stromal boundaries. (**D**) Plots of the difference between tip domain edge and stromal boundary segment orientation, mid-point position, and length. (**E**) Example Voronoi cells used to define tip domain boundaries shown overlaid on confocal micrographs. (**F-I**) *Left*, UB tip positions overlaid on height maps of representative kidney surfaces over E14-E17. *Right*, Voronoi diagrams tracing ‘tip domains’ overlaid on micrographs of combined ECAD and SIX2 immunofluorescence. Voronoi cells are colored by coordination number (# neighbors). (**J**) Frequency plot of tip domain coordination number (for (**F**)-(**I**), *n* = 5, 6, 4, 4 kidneys and 162, 353, 535, and 892 tip domains in E14, E15, E16, and E17 categories, respectively). (**K**) Average and standard deviation (s.d.) of coordination number distributions. (**L**) Sum of defect charge (i.e. net decrease in coordination number from *z* = 6) plotted against integrated Gaussian curvature for patches of kidney surface (units of disclinations).

We began by annotating UB tip positions on kidneys over embryonic days E14-E17. This allowed us to extract the predicted lattice boundaries of each tip ‘domain’ (all UB epithelium, cap mesenchyme, and stroma closer to a given tip than to a neighboring one) using a Voronoi tessellation approach (**Fig. 1C**). The Voronoi edges appeared to carry physical meaning, since they substantially overlapped stromal boundaries between domains. To quantify this, we annotated stromal boundary segments and found only small mismatches in their orientations (5.5 ± 4.4°, mean ± s.d., *n* = 64 segments), mid-point positions (5.8 ± 3.0 μm), and lengths (6.8 ± 4.7 μm) relative to the corresponding Voronoi edges (**Fig. 1D,E**). A similar correspondence has been noted in 2D foams and confluent cell monolayers ^30,31^, implying that the size and shape of tip domains are restricted by the presence of neighboring ones. These data show that tip domains can be modeled as tessellated polygonal sub-units with Voronoi boundaries equivalent to stromal boundaries.

We next annotated tip domains according to the number of neighbors they contact (the coordination number *z*), which is related to their ‘topological charge’ *s* = 6 - *z* (**Fig. 1E**). We then separately reconstructed the associated kidney surfaces from 3D confocal image stacks to compute their curvature maps (**Fig. S1**). Overlays of these data are shown in **Fig. 1F-I**. Tip coordination numbers varied between 4 and 8, and tip patterns qualitatively transitioned from a disordered state at E14-E15 to more visually apparent local order at E16-E17 (see also ^16^). The median coordination number of tips was *z* = 6, with a bias toward lower coordination numbers at earlier developmental times. This is consistent with tip patterns adopting energetically favorable pentagon disclinations (i.e. tips with *z* = 5 neighbors instead of 6) in younger kidneys with higher curvature (**Fig. 1J,K**). At later times, the coordination number distribution narrowed and approached a mean of 6, consistent with predictions for particle packings on flat surfaces (**Fig. 1K**). The theory of curved crystals suggests that the amount of bias toward pentagon dislocations will increase linearly with curvature (**Note S2**). Indeed, we find substantial agreement with the prediction of a bias toward pentagonal tip domains for younger kidneys with greater curvature per confocal region of interest (**Fig. 1L**). While variation in tip domain size and shape appears to contribute to the majority of excess dislocations in the kidney case, we also see evidence of grain boundary scars and clusters of defects of alternating charge (**Fig. 1F-I**). These data show that tip packing geometry at the kidney surface is partially determined by topological limitations imposed by kidney curvature.

### Crowding of ureteric bud tip domains creates crystal-like packing patterns and a mechanical rigidity transition

Beyond the contribution of curvature to the geometric microenvironment of UB tip domains, we sought to understand the impact of crowding among neighboring mesenchyme cap niches, each bordered by intervening layers of stroma. During early branching morphogenesis, caps and stroma are both fluid-like on a developmental timescale ^6,32^ (**Note S3**). However, together with an excess rate of new tip formation the stroma thins over E14-E18, which causes niches to pack closer together, conform in shape because of the confinement imposed by neighboring niches, and locally align with each other ^16^. Similar geometric features are observed in confined packings of soft elastic spheres ^28^ or closely packed droplets in dense two-phase emulsions such as foams ^33–35^. Since the degree of crowding can create abrupt changes in the geometric and mechanical properties of these systems, we wondered if similar effects could occur during niche packing. Due to the presence of stroma between caps, tip domain collectives have features of both non-confluent (density-dependent) and confluent (density-independent) packing. While hybrid models are emerging ^36^, we apply both areas of theory separately to gain insight on packing mechanics. One phenomenon in density-dependent packing is a rigidity (jamming) transition that occurs when droplets crowd together beyond a critical volume fraction *ϕ*_*c*_ and transition from zero to some finite yield stress (i.e. from fluid-like to solid-like behavior) ^37,38^. For kidney caps, we determined *ϕ* on a 2D basis as the ratio of cap area to total area. This area fraction *ϕ* exceeded those for 2D body-centered (square) packing of circles and for random close packing (which defines *ϕ*_*c*_ ^37,39,40^), even approaching that for 2D hexagonal close packing (hcp) after ∼E15 (**Fig. 2A**). This predicts a jamming transition, such that the ‘nephrogenic zone’ as a whole is predicted to be solid-like and therefore capable of imposing mechanical stress on newly formed niches during tip duplication after ∼E15.

**Fig. 2:**
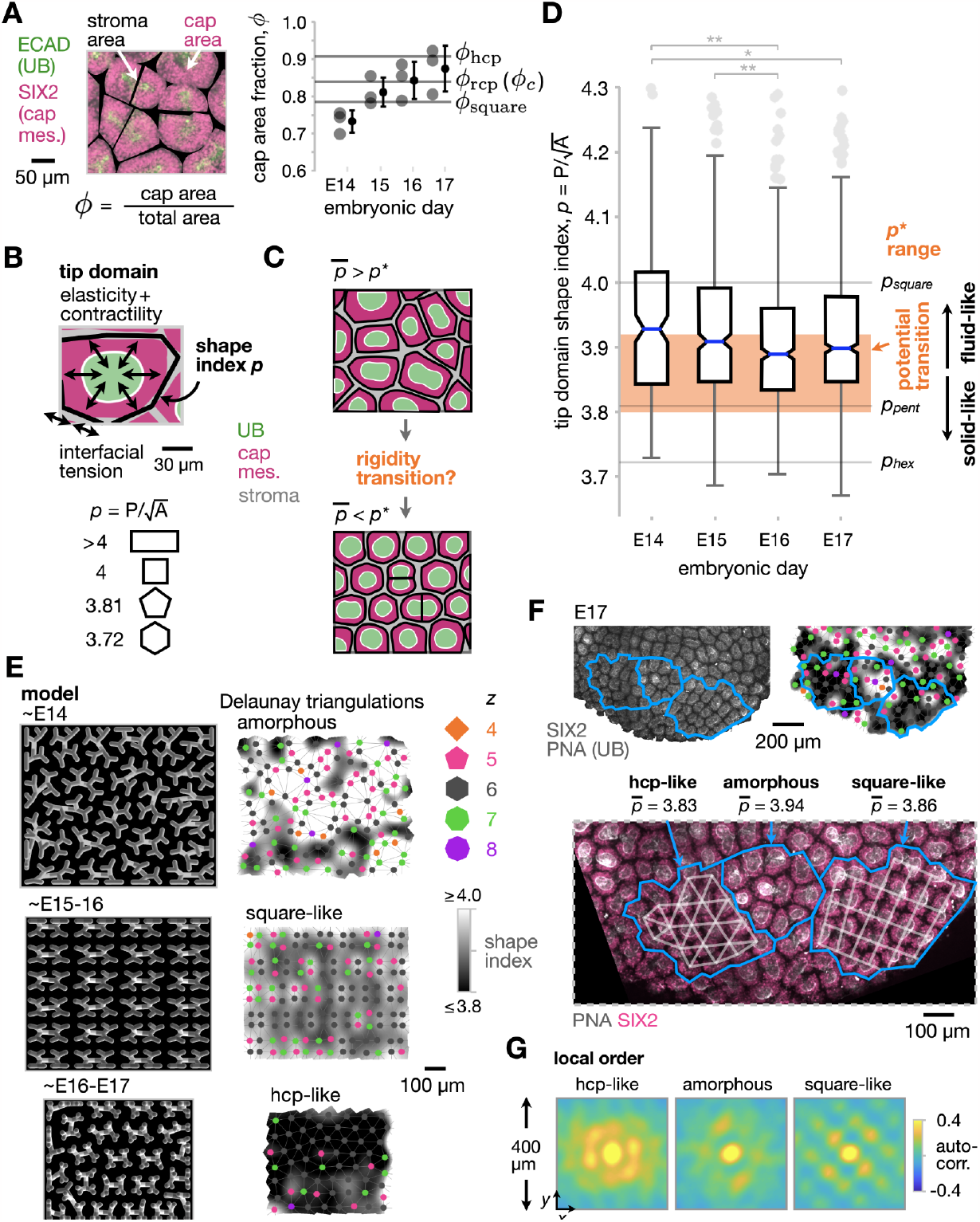
Close packing of ureteric bud tip domains predicts a transition to rigidity and emergence of semi-crystallinity over developmental time. (**A**) *Left*, Segmented confocal micrograph indicating the area fraction *ϕ* of cap mesenchyme. *Right*, Plot of *ϕ* over developmental time relative to 2D square packing, random close packing (rcp), and hexagonal close packing (hcp) of circles (mean ± S.D., *n* = 3 kidneys per embryonic day, average over > 12 tip domains per kidney). (**B**) Definition of shape index of tip domains *p* from Voronoi cell perimeter (P) and area (A). (**C**) Schematic of density-independent stiffening or rigidity transition when median shape index 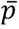 drops into a critical range. (**D**) Box plots of tip domain shape index distributions over time, relative to those for squares, pentagons (*pent*), and hexagons (*hex*), and to *p\** = 3.8-3.92 (*n* = 5, 6, 4, 4 kidneys and 177, 391, 617, and 987 tip domains in E14-17 categories, respectively. One-way ANOVA, Tukey’s test: *p < 0.05, **p < 0.01, ***p < 0.001). (**E**) *Left column*, Representative physical model output for tubule packing regimes characteristic of the indicated developmental ranges, previously reported in ^16^. *Right column*, overlays of Delauney triangulations indicating coordination number of tip domains on spatial heatmaps of shape index, showing reducing shape index as tip patterns transition from amorphous to square-like to hcp-like regimes. (**F**) Representative case study of amorphous and crystalline phases of tip domain packing at E17. *Top*, Confocal micrograph of tip domains and shape index heatmap. Example regions of different packing regimes are outlined in blue. *Bottom*, Inset of these regions showing lattice lines and decreasing median shape index 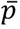 from amorphous to square-like to hcp-like packing. (**G**) Study of positional order within regions in (**F**) by spatial autocorrelation.

Although *ϕ* predicts density-dependent jamming of niches, the underlying theory neglects potentially important parameters ^37,41^. These parameters include collective cell elasticity and active contractility (embryonic kidney cortex explants shrink in the hours after cutting ^16^), and extra contributions to interfacial tension at cap-stroma boundaries (tissue layers are adhered through cell-cell junctions and/or cell-extracellular matrix interfaces). We therefore made some further predictions using density-independent rigidity theory created for the high packing fraction regime that does consider these parameters. Since niches pack side-to-side in one layer, we turned to a family of 2D vertex and Voronoi models that consider rigidity transitions in tissues organized into tessellated polygonal sub-units having elasticity, contractility and interfacial tension ^42–49^ (see **Note S4**). For any sub-unit with these properties such as individual cells or tip domains, the models predict significant stiffening or a sharp transition to rigidity when the median 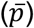 of a geometric parameter of their boundaries called the shape index *p* drops below a critical value *p\** in the range of 3.8-3.92 (**Fig. 2B,C, Note S4)** ^45,46^. For sharp transitions, when 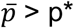 there is no energy barrier to changes in packing configuration, conferring fluid-like mechanics. However, when 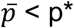 the configuration of polygons has both bulk and shear stiffnesses, conferring solid-like mechanics ^43^. To apply the model to the kidney, we measured the shape indices of tip domains over E14-E17, finding that the median shape index significantly decreased with developmental time, dropping into the critical range below 3.92 after ∼E15 (**Fig. 2D**). Despite a relatively small effect size, the change in shape index here is in a similar range to that previously associated with a rigidity transition in airway epithelial tissues *in vitro* ^50^. This result suggested that tip domain collectives may become stiffer and less viscous after E15, consistent with the time at which crystal-like locally ordered regions began to appear in **Fig. 1**. We next sought to connect these concepts and then to validate and quantify any rigidity transition using micromechanical experiments.

To draw a connection between shape index and our previous geometric model of tubule family packing ^16^, we produced heat maps of tip domain shape index for three characteristic regimes of tip packing. These three regimes, which at the tip domain level can be qualitatively referred to as amorphous, square-like, and hcp-like respectively, mirror the reduction in tip domain shape index from E14-E17 (**Fig. 2E**). In reality, tip packing does not perfectly adhere to any one regime at a snapshot in time (**Fig. 2F**). This could be caused by disruption of packed regions due to asynchronous tip duplication that would displace neighbors (**Movie S2A**) and create heterogeneity (polydispersity) in the relative size and shape of tip domains (quantified later in **Fig. 4**). This means that our previous model predicts the stage at which a given packing phase has the potential to exist, but a mixture of phases arises in practice. For example, at E17 we observe regions that exhibit amorphous, square-like, or hcp-like packing geometry. These regions persist spatially for 2-3 tip spacings in *xy*, as revealed by spatial autocorrelation analysis showing four-fold and six-fold rotational symmetry for square-like and hcp-like regions respectively (**Fig. 2G**). Overall, these data show that tip domain packing at the kidney surface is semi-crystalline with an increased prevalence of square- and hcp-packed regions at later developmental times.

We next wondered if our predictions from rigidity theory and the increase in crystalline order of tip domains over time would reveal stiffening or a sharp rigidity transition after E15 on the length-scale at which tip domains interact. We hypothesized that this would manifest both temporally and spatially - firstly that the overall decrease in *p* and increase in tip domain crystallinity over time would increase tissue stiffness, and that tip domains in regions of higher crystalline order (lower local average *p*) should show higher stiffness. Taking advantage of the accessibility of tip domains at the kidney surface, we employed surface microindentation and Brillouin microscopy to quantify temporal and spatial changes in tissue mechanics, respectively.

We tackled the time-dependent aspect using surface microindentation of freshly explanted kidneys to quantify elastic modulus (stiffness) and viscoelasticity over E15-E17 using a 254 μm cylindrical indenter (equivalent to the width of ∼3 tip domains, **Fig. 3A**, see **Methods**) ^51^. We measured the applied force recorded during indentation of the kidney surface by ∼30 μm, and paused indentation to capture the time dependence of force during subsequent tissue relaxation (**Fig. 3B**). These measurements revealed a significant increase in kidney surface stiffness local to tip domains over E15-17 (**Fig. 3C,D**). Secondly, they showed an increase in the time-scale of force relaxation over E15-E17, perhaps due to a slowing in passive remodeling of cell collectives and extracellular matrix ^51^. Thirdly, the measurements showed a decrease in viscous relaxation over E15-E17, perhaps caused by slowing of active tissue remodeling. Together, these data reveal marked increases in stiffness and decreases in passive and active tissue remodeling over E15-E17, the same time period over which our tip domain shape index analysis predicted tissue stiffening (if not a sharp fluid-like to solid-like rigidity transition) (**Fig. 2D, Fig. 3E**).

**Fig. 3:**
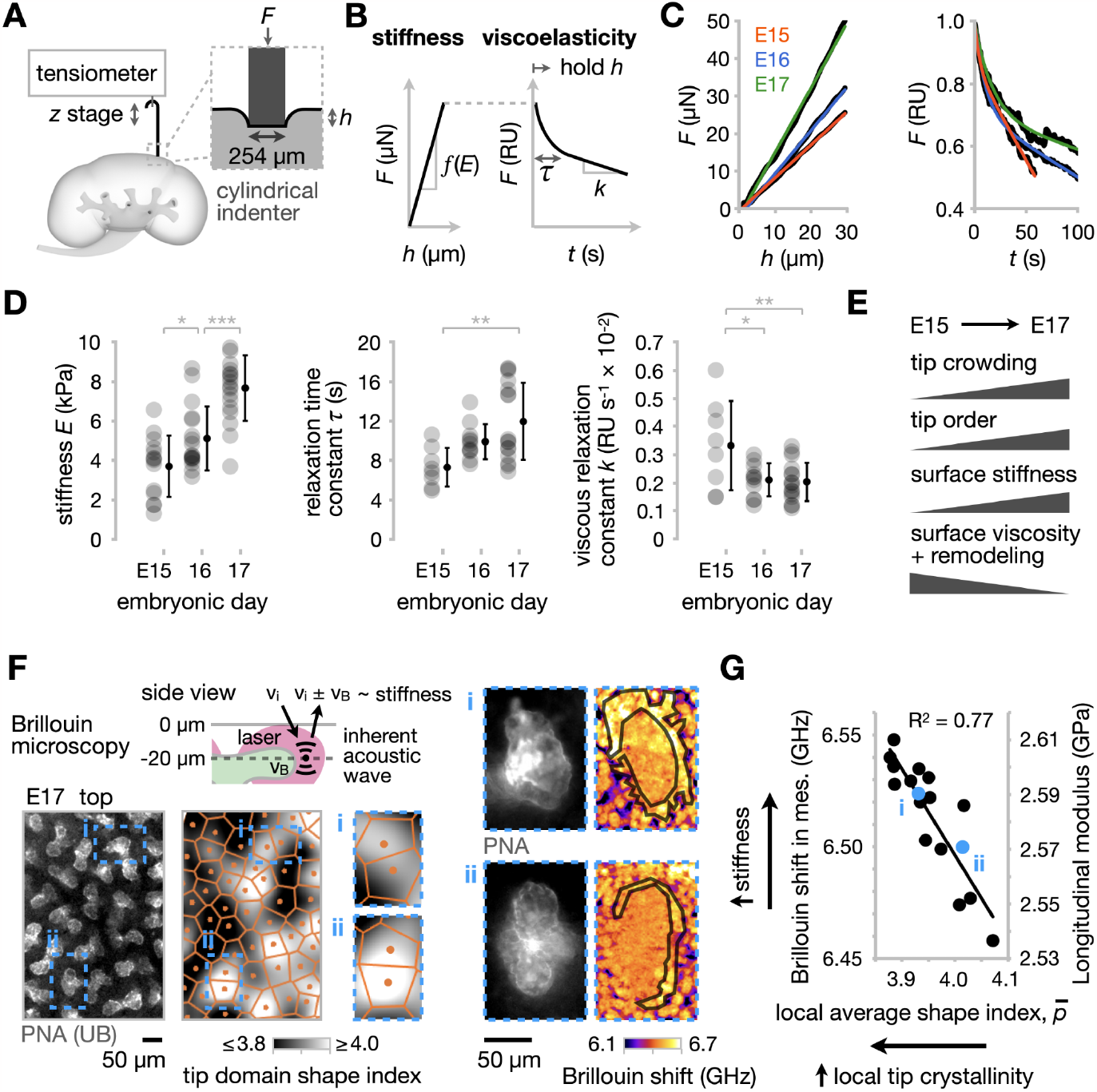
Crowding of ureteric bud tip domains stiffens the nephrogenic niche over time, especially in close-packed regions. (**A**) Schematic of surface microindentation using a cylindrical indenter that captures kidney viscoelastic properties on the length scale of ∼3 tip domains for *h* ∼ 30 μm. (**B**) Schematic of typical data output and key parameters during indentation (*left*), and during subsequent recording of force at fixed final indentation depth (*right*). (**C**) Representative indentation force vs. indentation depth, and force relaxation curves for E15-E17 kidneys. (**D**) Stiffness, relaxation time *τ*, and viscous relaxation constant *k* plots over E15-E17 (mean ± S.D., *n* > 14 kidneys per embryonic day for stiffness and > 8 for *τ* and *k*, one measurement per kidney. One-way ANOVA, Tukey’s test: *p < 0.05, **p < 0.01, ***p < 0.001). (**E**) Summary of changes in UB tip geometry and kidney surface mechanics over E15-17. (**F**) *Top*, Schematic of Brillouin microscopy, which quantifies an optical frequency shift that depends on tissue stiffness. *Bottom*, live confocal micrograph of PNA-labeled UB tips, Voronoi diagram of tip domains with heatmap of shape index, and zoomed insets of two example domains. *Right*, Insets of the two example domains with PNA fluorescence and Brillouin shift images. Cap mesenchyme cells local to tips are outlined. (**G**) Mean Brillouin shift and corresponding estimated longitudinal modulus in cap mesenchyme cells vs. average tip domain shape index in their local neighborhood of ∼3 tips (*n* = 17 tip domains across 4 E17 kidneys).

**Fig. 4:**
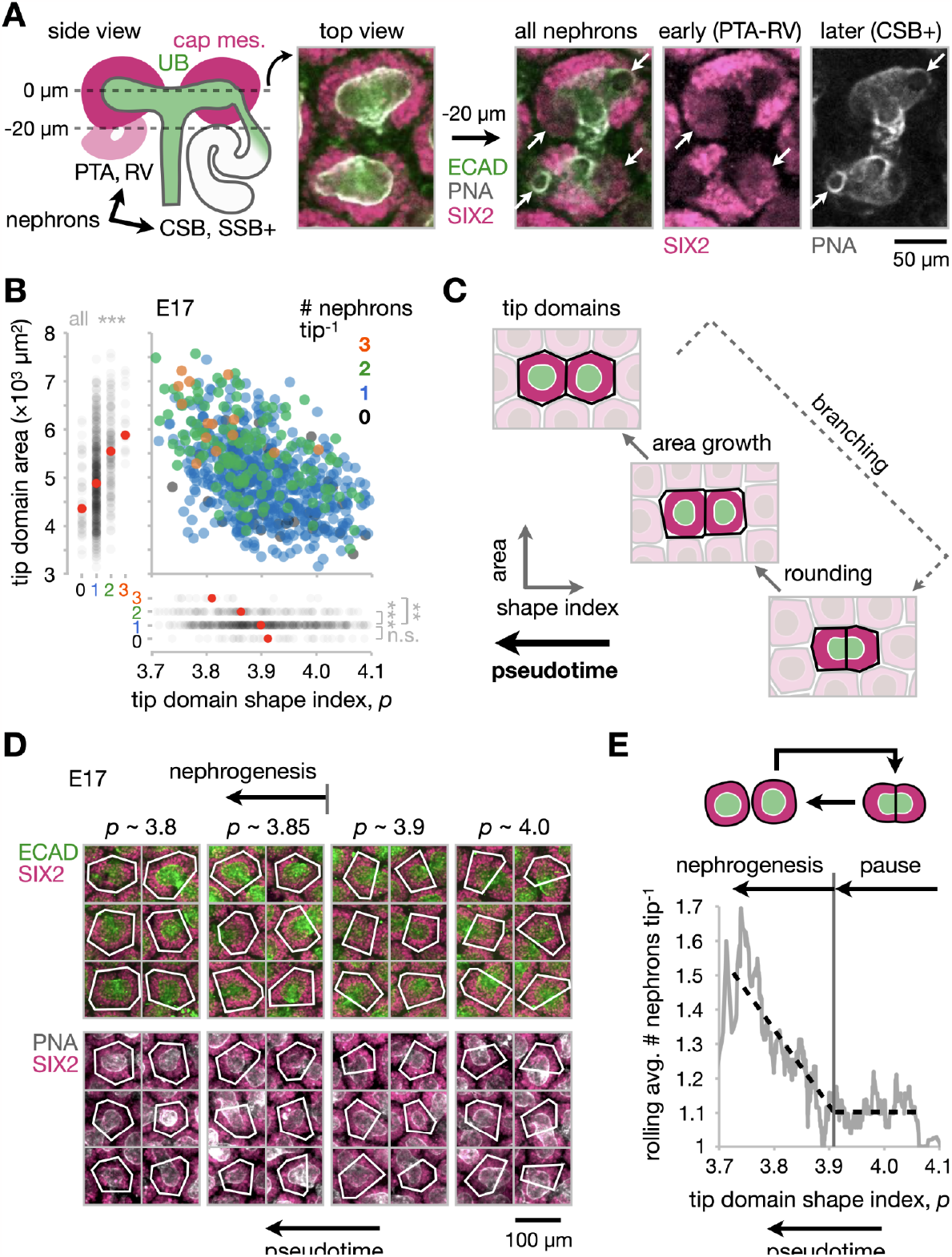
Nephrogenesis rate varies over a tip domain ‘life-cycle’ defined by shape index. (**A**) *Left*, Schematic showing side (*xz*) view of a pair of UB tips associated with early stage nephrons (PTA, pre-tubular aggregate; RV, renal vesicle) and later stage nephrons (CSB, comma shaped body; SSB, S-shaped body). *Right*, Confocal immunofluorescence micrographs at the mid-plane of UB tips and 20 μm deeper into an E17 kidney. All nephrons associated with each tip can be annotated either as SIX2^+^ spheroids for early nephrons or from the attachment points of their connecting tubules for later nephrons (white circles at tips in PNA channel), see arrows. (**B**) Plot of tip domain area against shape index, with nephron number per tip in the color dimension (*n* = 694 tip domains across 4 E17 kidneys. One-way ANOVA, Tukey’s test: *p < 0.05, **p < 0.01, ***p < 0.001). Each axis is also plotted individually vs. nephron number per tip. (**C**) Schematic of UB tip life-cycle and definition of a pseudotime based on shape index of tip domains. (**D**) Example confocal micrographs of tip domains over a range in shape index. (**E**) Rolling average of nephron number per tip against shape index (window width = 50 points).

We next sought to validate the spatial aspect of our hypothesis using confocal Brillouin microscopy ^52,53^, a label-free optical technique that has been previously used to quantify the embryonic tissue mechanics of the spinal cord, cornea, and fly ventral furrow during gastrulation ^54–56^. Brillouin microscopy measures the frequency shift (i.e. Brillouin shift) between incident and scattered light caused by the interaction of incoming photons with inherent (spontaneous) acoustic waves in biological materials ^57^ (**Fig. 3F**). The Brillouin shift scales with the high-frequency longitudinal modulus, a uniaxial stress-to-strain ratio that is proportional to the more widely known quasi-static Young’s modulus *E* (‘stiffness’) in cells and tissues over a physiological range relevant to embryonic development ^53,58^. We acquired 2D Brillouin images of randomly chosen E17 tip domains at planes approximately bisecting tubule tips and surrounding mesenchyme (**Fig. 3F**). We found that the mean Brillouin shift in cap mesenchyme cells and the average shape index in a local neighborhood of ∼3 tips were correlated (**Fig 3G**); specifically that the longitudinal modulus of the cap mesenchyme is significantly higher for tip domains predicted to reside in more closely packed and solid-like regions with crystalline order (**Fig. 2D-G**). Although accurately translating the high-frequency Brillouin modulus to the quasi-static Young’s modulus requires careful calibration specific to a given sample, we can approximate the relative change in Young’s modulus based on an empirical log-log linear relationship established using cells ^57^. For a Brillouin shift increase of ∼0.1 GHz, the relative increase in Young’s modulus is ∼46%. These data show that the stiffness of nephrogenic niches varies substantially over the kidney surface and is significantly higher for tips in regions with more crystalline packing.

An increase in kidney surface stiffness and reduction in viscosity would increase the mechanical resistance to new tip domains forming, expanding, and displacing close-packed neighbors, creating residual stress (**Movie S2B**). This would imply that tip domains see cyclical swings in their mechanical microenvironment on a similar timescale to tip duplication and nephron formation events. To establish this, we must first define a tip domain ‘life-cycle’ between duplication events and then assay mechanical stress in niches at different points in their life-cycles.

### Correcting for asynchronous branching by defining the tip domain life-cycle in terms of a common pseudotime

Establishing a life-cycle estimate for each niche would enable instantaneous stress measurements from different niches to be aligned in a common ‘pseudotime’ axis that would correct for asynchronous duplication of niches. Creating a pseudotime axis is necessary because it is not currently possible to perform live time-lapse imaging of UB tip domain branching in kidney explants while retaining the *in vivo* 3D architecture of the organ (without it becoming spheroidal or flattened) ^59–61^. An appropriate pseudotime axis would circumvent this by allowing us to estimate an averaged stress transient in niches from fixed time-point kidneys. We focused on E17 kidneys, reasoning that any periodic increase in local mechanical stress due to tip duplication would be amplified because of their higher surface stiffness and lower viscosity relative to earlier stages. Since nephrons are distributed among daughter tips when a tip divides ^4^, we reasoned that among those present at the same developmental time-point the tips bearing larger nephron numbers should be ‘older’ relative to their last branching event. Nephrons can be scored from 3D confocal immunofluorescence stacks by combining annotations of SIX2+ spheroids beneath UB tips (capturing pre-tubular aggregate and renal vesicle stages) with annotations of sites where connecting tubules from more mature nephrons connect to tips (capturing comma shaped body, S-shaped body, and later stages), **Fig. 4A**. First, we found a negative correlation between tip domain area and shape index, reflecting the proposed ‘life-cycle’ of tips. Specifically, recently divided tips bearing fewer nephrons have lower area and higher shape index, while older tips bearing more nephrons are larger with lower shape index (**Fig. 4B,C**). As a tip ages and gains nephrons, its domain grows in area and becomes more circular before the cycle repeats upon its next branching event. These data imply that the tip domain shape index can be used as the pseudotime dimension that corrects for the lack of synchronization of tip branching. In other words, tips in different locations can be aligned with respect to their progress through morphogenesis relative to their last duplication event (**Fig. 4D**). This enabled us to plot a rolling average of nephron number per tip against shape index. If committed nephron progenitors condense into discrete early nephrons at a fixed rate, we would not expect the nephrogenesis rate to depend on tip life-cycle. However, nephrogenesis appears to switch on as the shape index of tip domains drops below an intermediate value of *p* ∼ 3.91 and the area of tip domains begins to increase (**Fig. 4E, Fig. S2**). This means that nephrogenesis pauses as new tips push outward against neighboring tip domains and begin to round up, and resumes as tip domains then grow in area.

### Newly branching tip domains experience mechanical stress that dissipates over their life-cycle

Having defined a life-cycle for tip domains, we went back to test if tip domains see cyclical swings in their mechanical microenvironments that synchronize with their life-cycles. Specifically, we predict that this would be caused by branching tips overcoming mechanical resistance while displacing close-packed neighbors, creating transient stress. To study this, we first drew on force inference methods developed from cell-scale vertex modeling to investigate the spatial distribution of mechanical stress in tip domain collectives ^62–64^. We reasoned that force inference could be extended to the tissue level because both cells and tip domains can be modeled by networks of interfaces (cell junctions / stromal junctions) that transmit mechanical stress between neighbors. Parallel (interfacial tension) and perpendicular (hydrostatic pressure, elasticity, and contractility) forces create observed variations in interface edge lengths and angles at vertices. Mechanical stresses can be determined from these data without assuming the biophysical origin or spatial localization of forces ^62–65^ (**Fig. 5A, Note S5**). Force inference has been successfully validated by laser ablation experiments across several model organisms, tissue types, and length-scales (from several-cell to highly multicellular tissues) ^64^. We performed Bayesian force inference from E17 kidney surface projections using tip domain Voronoi edges ^66^ that outline stromal interfaces between domains (**Fig. 1C,D**). This produced a spatial map of the inferred stress tensor, which yields major and minor principal stress axes and relative magnitudes that give relative isotropic and anisotropic (deviatoric) stresses (**Fig. 5B**). We plotted these components against the shape index of tip domains to show that as tips age relative to their last duplication event (i.e. as shape index decreases), they see a marginal increase in inferred isotropic stress, and a ∼30% decrease in anisotropic stress across stromal interfaces (**Fig. 5C**). This indeed predicts a rhythmic mechanical microenvironment synchronized with the tip life-cycle, and in particular, that mechanical tension immediately precedes the nephrogenic phase (**Fig. 5C, Fig. 4E**).

**Fig. 5:**
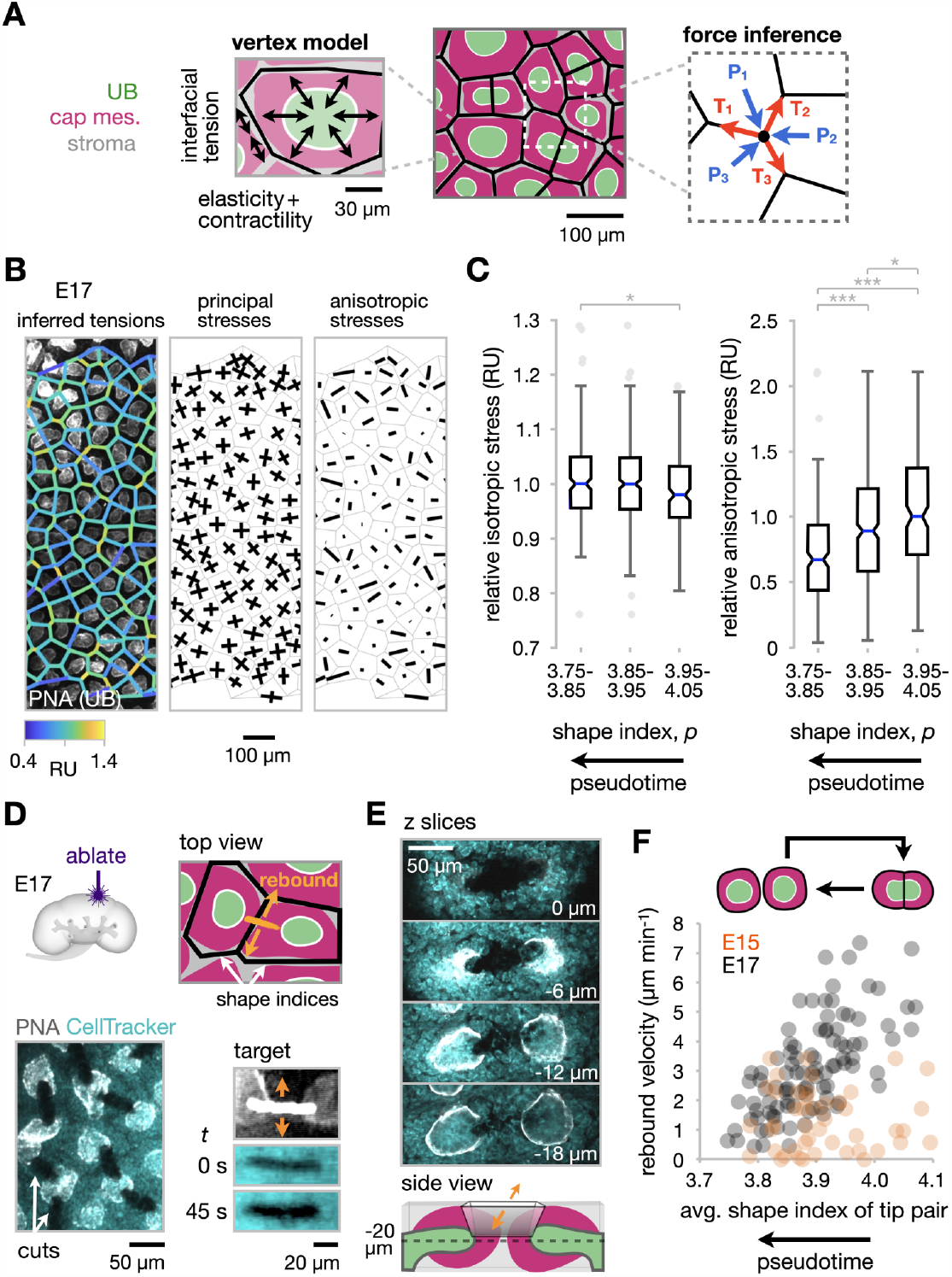
Newly established tip domains experience mechanical tension that decreases over each life-cycle. (**A**) Schematic of inferred tension (T) and pressure (P) contributions to the force balance at an example node in a vertex model representation of tip domains. (**B**) Example force inference output from tip domain Voronoi geometry as a spatial map overlaid on confocal fluorescence micrograph of E17 kidney surface, along with relative principal and anisotropic stress vectors. (**C**) Box plots of isotropic and anisotropic stress for tip domains, relative to the median of the highest stress category in each plot (*n* = 4 kidneys; 195, 319, and 164 tip domains in low, medium, and high shape index categories, respectively. One-way ANOVA, Tukey’s test: *p < 0.05, **p < 0.01, ***p < 0.001). (**D**) *Top*, Schematic of laser microdissection ablation of cap mesenchyme and stroma orthogonal to boundaries between neighboring tip domains at E17. *Bottom*, Confocal micrograph showing example cuts after fixation, cut location prior to ablation by widefield fluorescence, and rebound of the cut in the 45 s after ablation. (**E**) Sequential z-slices in a confocal stack after fixation, showing penetration of a cut ∼20 μm through the inter-tip region (cell integrity disrupted down to ∼40 μm). (**F**) Plot of cut rebound velocity (change in cut width in 45 s) after ablation against the average shape index of the tip domains immediately adjacent to each cut (*n* = 45 and 99 cuts pooled across 5 and 7 kidneys for E15 and E17 respectively).

We next sought to validate inferred stresses local to tip domains experimentally. We used laser microdissection of E17 kidneys shortly after explanting to create slot-like ablations spanning neighboring cap mesenchyme populations and orthogonal to the stromal interfaces between pairs of UB tips (**Fig. 5D, Fig. S4, Movie S3**). Laser ablation is thought to most closely reflect anisotropic stress at cutting sites ^63^, such that the rebound (opening) velocity of cut edges scales with local tension ^64^. Ablation successfully induced rebound at cut sites (**Fig. 5D, Fig. S4**), and cuts penetrated ∼40 μm in depth, sufficient to sever the mesenchyme and stroma between tips over their full *z* extent (**Fig. 5E**). Validating the force inference data, the rebound velocity of cuts significantly increased with the average shape index of tip domains adjacent to each cut at E17 (**Fig. 5F, Fig. S3, Fig. S4**). These data show that anisotropic stress local to the cap mesenchyme is highest for newly duplicated tips with higher shape index and lowest for older tips with lower shape index. This held for E17 kidneys but not for E15 kidneys prior to the rigidity transition, where residual stresses would be predicted to dissipate and not be transmitted among domains (**Fig. 5F, Fig. S3**). The data validate the stress inference model and our earlier notion that in kidneys that have crossed the rigidity transition newly formed tip domains are subjected to mechanical stress as they overcome mechanical resistance while branching and displacing close-packed neighbors. This tension then dissipates prior to the next duplication cycle (**Movie S2B**).

## Discussion

Gaining a fundamental understanding of tissue-wide coordination between ureteric bud, nephron, and stromal morphogenesis may lead to control strategies that correct variability in nephron endowment in kidneys and in kidney organoid models ^22,67,68^. Organoids constitute both a platform to clarify the fundamentals of nephrogenesis and a potential vehicle for augmentation of adult kidney function. However, nephrons form in a single synchronous pulse in iPSC-derived kidney organoids rather than in an ongoing, periodic manner ^69,70^. Replicating the latter in organoids is a significant opportunity since successive rounds of nephrogenesis is crucial to achieving high nephron density in *vivo*. However, not enough is known about periodic niche regulation. Several biochemical pathways coordinate non-autonomous signaling between the UB, cap mesenchyme, and stromal compartments. These include Wnt/β-catenin, BMP/MAPK and SMAD, planar cell polarity, FGF, GDNF/Ret, retinoic acid and others ^7,8,71–80^. Among these, an appropriate tuning of ligand concentration, presentation geometry/transport, and cell perception sensitivity is thought to set an optimal ratio of cell renewal (vs. priming) vs. differentiation events within each compartment ^8,68,81,82^. These features would change over the developmental span to generate an initial period of increasing nephrogenesis rate relative to branching rate, followed by a period with constant rate, ending with a pulse at cessation of nephrogenesis ^1,4,68,81,83^. This perspective is satisfying when averaged over niches, but does not explain how e.g. differentiation cues are regulated in response to radical changes in niche geometry caused by growth and splitting cycles synchronized with UB branching events ^4,16^. Therefore, an emerging area of interest is to determine potential geometric and mechanical factors that may add context-dependent cues to cell decision-making in and adjacent to nephron progenitor niches. These include the mutual inhibition of UB tip branching based on local crowding ^15^, the interplay between nephron progenitor number per tip and nephrogenesis rate ^22^, and niche shape- and cell migration-dependence of progenitor exposure to pro-renewal vs. -differentiation ligands ^9,84^.

Supporting these efforts, our work offers fundamental characterization of nephrogenic niche geometry and mechanics over the UB tip branching life-cycle. The results suggest that topological requirements imposed by decreasing curvature of the kidney and closer packing of tip domains forces them into semi-crystalline organization after ∼E15 (**Fig. 6A**). Two bodies of theory predict density-dependent and -independent contributions to a rigidity transition in tip domains that coincide with this. We note that flattening of the kidney surface as it grows could also reinforce the transition by increasing the critical shape index at which rigidification occurs ^85,86^. The predicted rigidity transition coincides with an experimentally measured increase in solid-like properties at the kidney surface, creating the potential for residual stresses to exist. Physical resistance to tip duplication as they displace jammed neighbors would mean that each tip domain experiences rhythmic temporal waves in mechanical stress (**Fig. 6B**). Indeed, we find that mechanical stresses local to tip domains correlate with their life-cycle at E17 but not prior to the rigidity transition at ∼E15. Together with the observation that UB tips progressively over-crowd the kidney surface ^16^, these data challenge a null hypothesis that new niches form as sufficient surface area becomes available. Rather, the data show that niches physically compete for occupancy at the surface after ∼E15, creating measurable mechanical stress local to each asynchronous branch event.

**Fig. 6:**
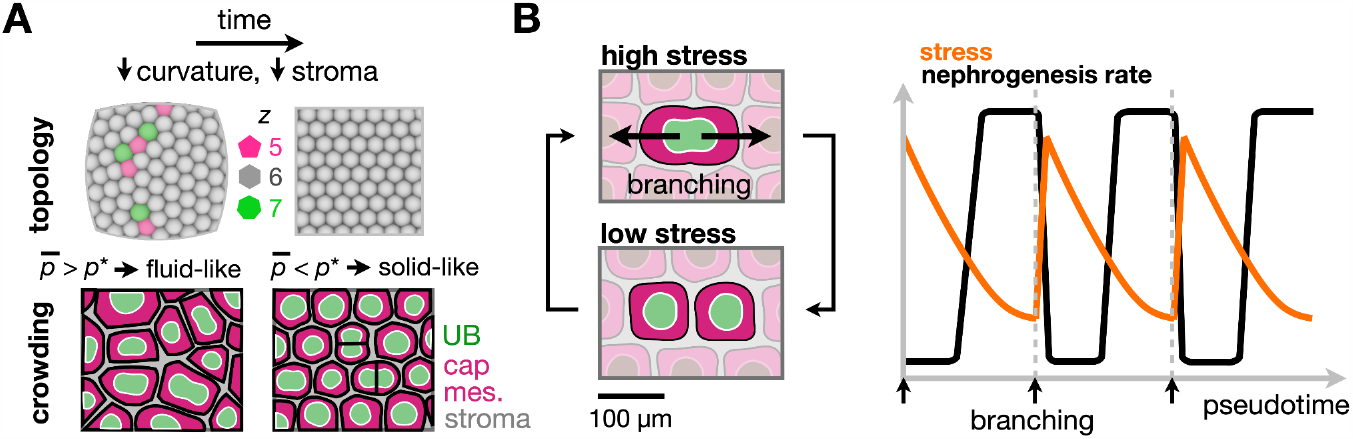
Model for curvature and crowding contributions to tip domain packing and its effects on mechanics in the nephrogenic niche. (**A**) Schematic summary of curvature and crowding contributions to tip domain geometry and rigidity transition. (**B**) Schematic summary of cyclical mechanical stress and nephrogenesis in the tip domain microenvironment associated with the tip duplication life-cycle.

Our data connecting niche geometry, mechanics, age since duplication, and nephrogenesis rate set up several areas for future investigation *in vivo* and in organoids. For example, correlations between niche age since duplication and cell renewal/differentiation pathway activity could be assayed in fixed kidneys. This may inspire new organoid engineering approaches seeking to maintain both naive niches and differentiating nephrons in the same organoid. Our data on rhythmic mechanical stress in niches raises a second follow-up - is stress in the UB, cap mesenchyme, and/or stromal compartments instructive to cell decision-making? This is motivated by observations including that the BMP/pSMAD and Wnt/β-catenin axes that prime and differentiate nephron progenitors are mechanosensitive in other contexts ^87–89^. Secondly, that nephron progenitor regulation is dependent on the mechanotransduction pathway member YAP ^90–93^ and on appropriate Rho/ROCK and non-muscle myosin II activity ^91,94,95^ (**Note S6**). This question could be tackled through mechanobiology studies in mouse kidney explants, primary cells, and reconstituted or iPSC-derived organoid systems. Together these efforts will lead to a broader understanding of connections between kidney geometry, mechanics, and niche signaling toward achieving greater engineering control over nephron formation in regenerative medicine applications.

## Acknowledgements

We would like to thank Hughes lab members, Zev Gartner, Celeste Nelson, and Melissa Little for discussion and advice. We are grateful to Kate Bennett at the Molecular Pathology and Imaging Core, Gastroenterology Division, Penn Medicine for technical assistance with laser ablation studies; and David Li and Paul Janmey for access to and training on microindentation. This work was supported by: NIH F32 fellowship DK126385 (LSP), the Predoctoral Training Program in Developmental Biology T32HD083185 (JMV), NSF GRFP (CMP, JL), NIH NIGMS MIRA R35GM133380 (AJH), NIH NIDDK R01DK132296 (AJH), NSF CAREER award 2047271 (AJH).

## Author contributions

Conceptualization: JMV, LSP, TJC, AJH

Methodology: JMV, LSP, JL, AH, TJC, GHL, CMP, JZ, AJH

Software: JMV, TJC, JZ, AJH

Formal analysis: JMV, TJC, JZ, AJH

Investigation: JMV, LSP, JL, AH, TJC, GHL, CMP, CS, JZ, AJH

Writing - original draft: JZ, AJH

Writing - review and editing: JMV, LSP, JL, AH, TJC, GHL, CMP, JZ, AJH

Visualization: JMV, LSP, TJC, GHL, CS, JZ, AJH

Supervision: LSP, JZ, AJH

Project administration: JZ, AJH

## Data and materials sharing

All data necessary to evaluate conclusions of this study are presented in the paper and supporting information. In-house code and raw data are available from the authors upon request.

## Supporting information

### Supporting figures

**Fig. S1:**
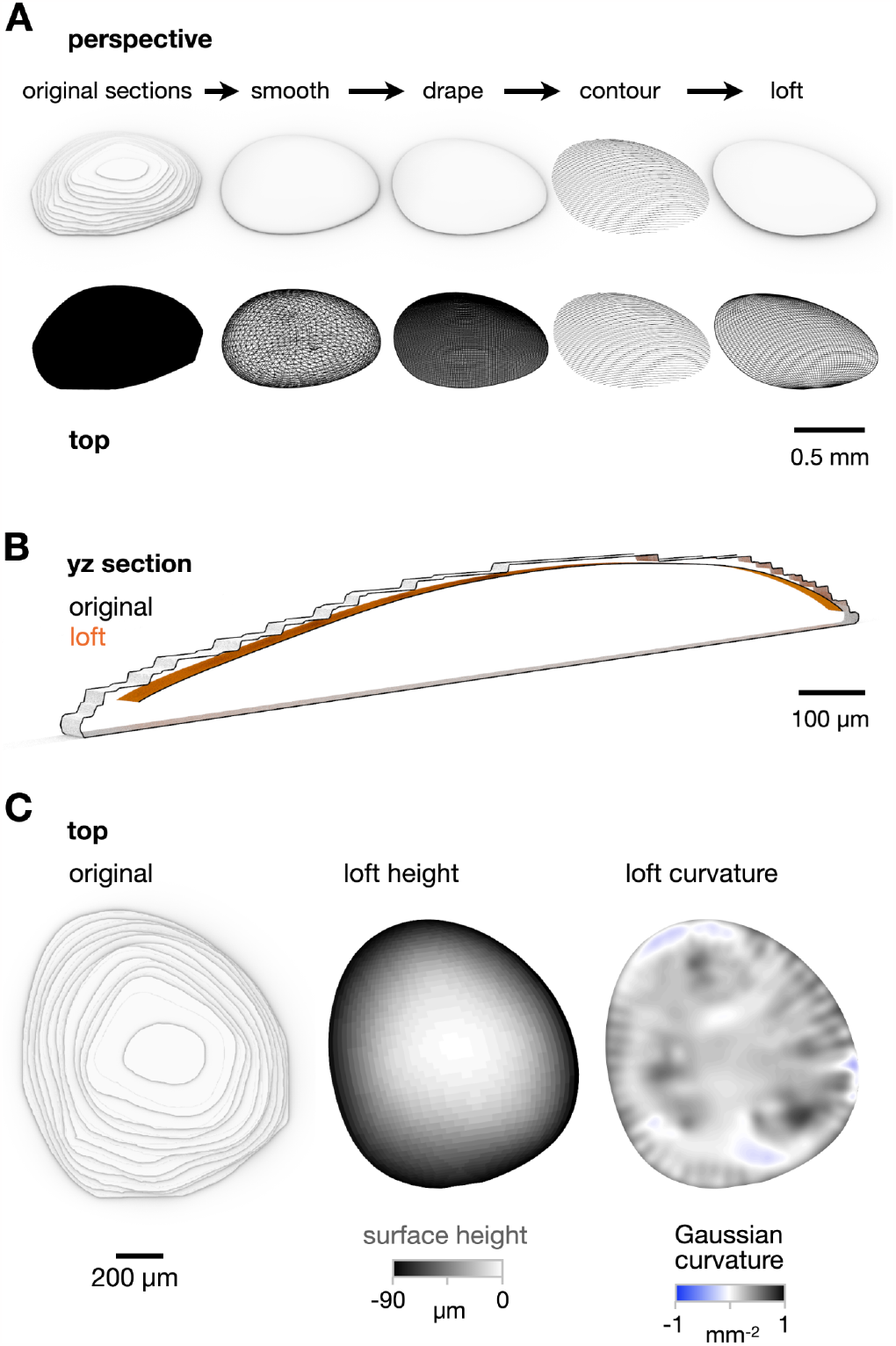
Kidney curvature analysis. (**A**) Perspective rendering (*top row*) and wireframes (*bottom row*) for example segmented E17 confocal stack and resulting surfaces generated after each processing step in Rhino (see **Methods**). (**B**) Midplane *yz* cross-sections of original segmented volume and loft surface after pre-processing steps, showing appropriate fit for subsequent curvature analysis. (**C**) Top (*xy*) view renderings of original volume, loft surface as a heatmap of height from MATLAB, and loft surface as a heatmap of Gaussian curvature from Rhino Grasshopper.

**Fig. S2:**
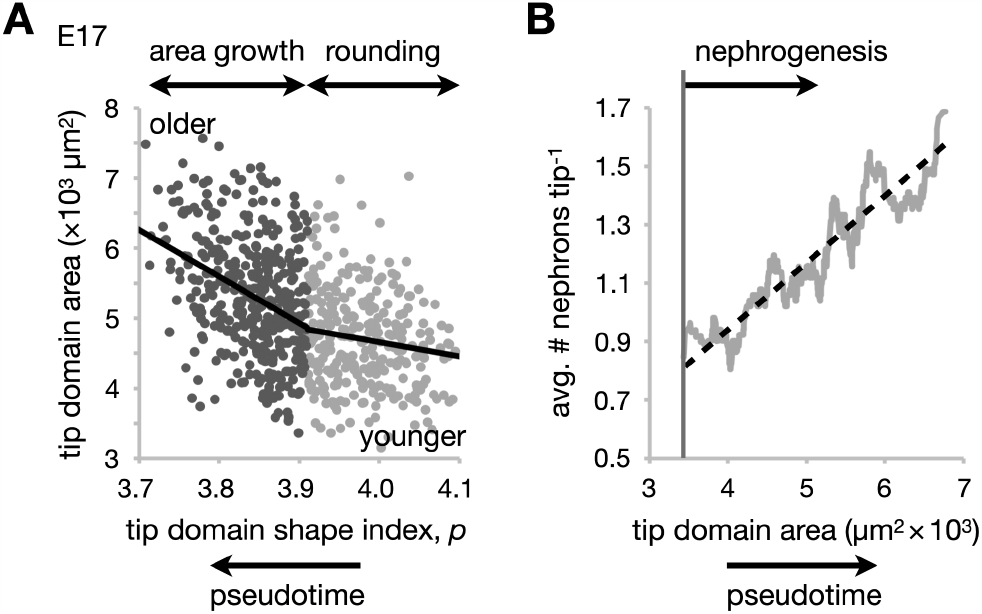
Tip domains transition from rounding-dominated to area growth-dominated regimes over tip life-cycle. (**A**) A version of **Fig. 4B** divided into two periods during the tip life-cycle before (right side) and after (left side) nephrogenesis begins (**Fig. 4E**). Note that younger tip domains with fewer nephrons primarily round up prior to the nephrogenesis phase, which coincides with a higher rate of tip domain area increase. (**B**) A version of **Fig. 4E** except plotted against tip domain area rather than shape index.

**Fig. S3:**
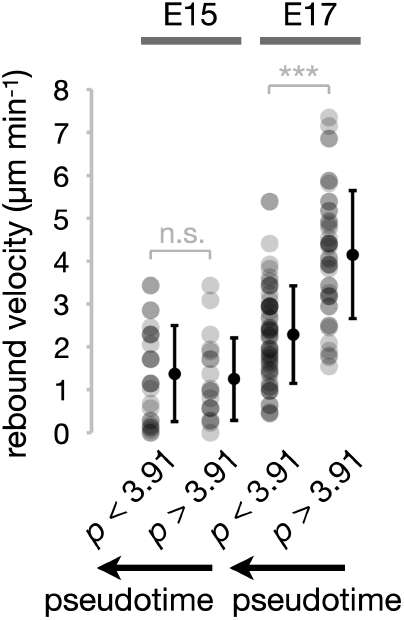
Newly established tip domains experience anisotropic stress that decreases over each life-cycle at E17 but not prior to the rigidity transition at E15. Plot of rebound velocity of cuts (change in cut width in 45 s) after ablation against the average shape index of the tip domains immediately adjacent to each cut, binned into low (*p* < 3.91) and high (*p* > 3.91) shape index groups (mean ± S.D., *n* = 26, 19, 56, and 43 cuts from left to right, pooled across 5 and 7 kidneys for E15 and E17 respectively. One-way ANOVA, Tukey’s test: *p < 0.05, **p < 0.01, ***p < 0.001).

**Fig S4:**
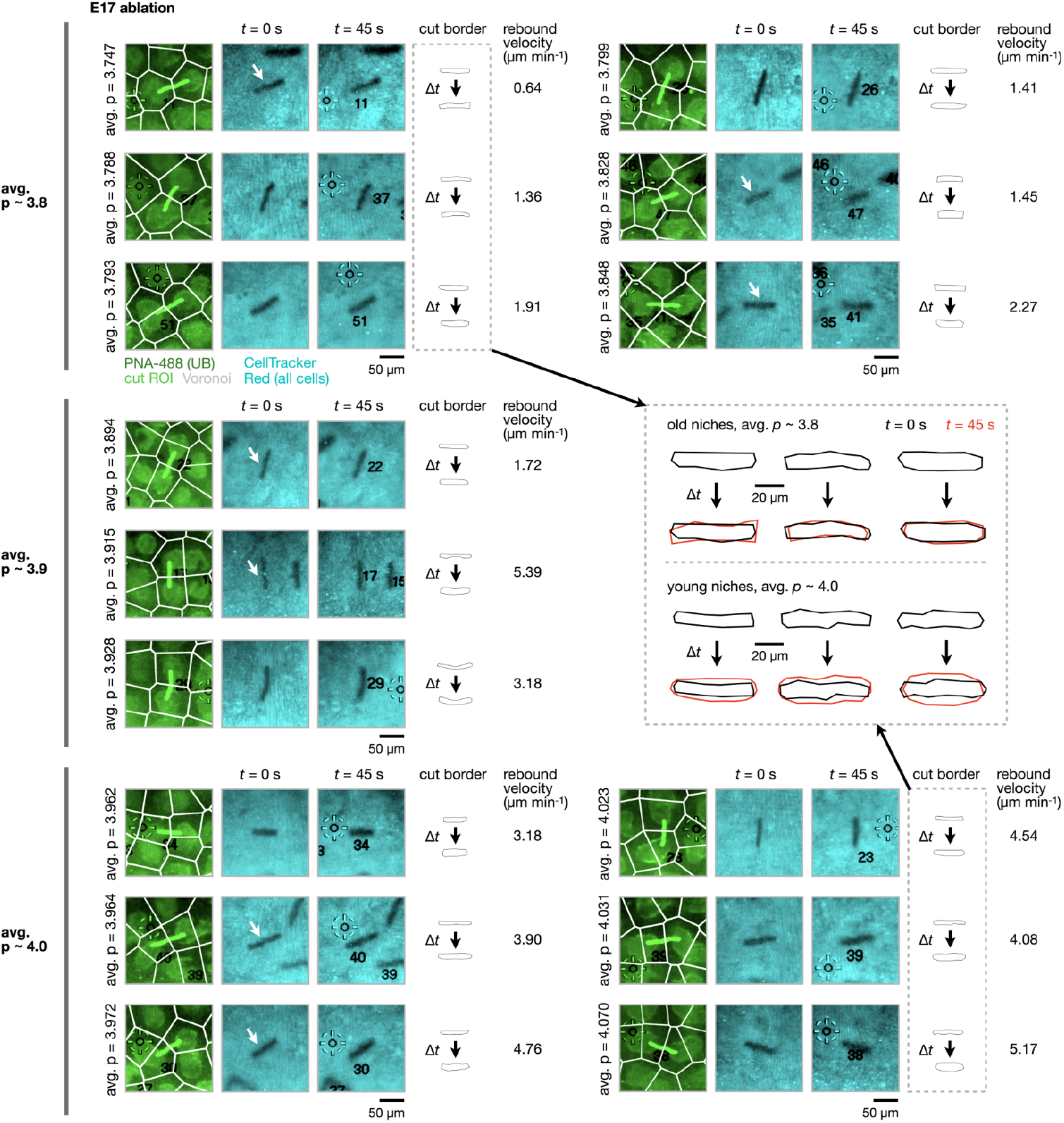
Laser ablation case studies over the niche life-cycle. Example laser ablation cases for 15 of the 99 cuts on E17 kidneys contributing to Fig. 5F. Representative cases are displayed for average shape index of the tip domains immediately adjacent to each cut in the vicinity of p ∼ 3.8, 3.9, and 4.0 to visually contrast cut opening magnitudes among these categories. Each case shows fluorescence micrographs of UB tips with cut ROIs and local Voronoi diagram overlaid, and kidney surface fluorescence immediately after cutting and after 45 s. Cut boundaries and associated rebound velocities are also shown. White arrows indicate the cut of interest if more than one cut appears in the field of view. *Inset*, zooming in to show boundaries for a representative subset of cuts, emphasizing higher rebound magnitudes for cuts made between younger niches having higher average shape index that are earlier in their branch life-cycles.

### Supporting movies

**Movie S1: Coordination of ureteric bud tip branching morphogenesis and nephron formation**. Side-view schematic depicting two cycles of UB tip duplication. UB tubules (green), cap mesenchyme (magenta), early nephrons (white). UB tips branch at the surface of the kidney and are surrounded by cap mesenchyme cell populations that repel each other. Early nephrons are induced at ‘armpit’ regions of tubules. Note that nephrons are distributed among daughter tips during each duplication cycle. The movie represents the E15-P2 period during which tip duplication rate matches nephron formation rate ^4^. Box inset represents the ‘self-similar’ nature of branching.

**Movie S2: Asynchronous tip duplication disrupts crystal-packed regions of tip domains *in silico***. (**A**) Conceptual movie of model tip domain pairs represented by sphere pairs repelling other pairs at an example E15 mouse kidney surface mesh. Tip domain pairs are added at random locations as the mesh grows isotropically such that tip domain number increases proportional to mesh surface area to the power 1.3 (ref. ^16^). Spatial parameters approximately reflect ∼E15-E18. Note the emergence of square- and hexagonal close-packed regions that are disrupted spatially and temporally by tip duplication events. (**B**) Inset of another simulation instance over the latter portion of (**A**) in which sphere pairs are shaded by relative local stress (proportional to elastic strain between pairs). Note local stress increases upon spatially random addition of new tip domain pairs.

**Movie S3: Laser ablation causes rebound of cap mesenchyme and stroma between pairs of ureteric bud tips**. Timelapse of E17 mouse embryonic kidney by widefield microscopy during laser microdissection (see **Methods**). The timelapse is played at ∼10x real-time. UB tips are shown firstly by Alexa Fluor 488-PNA fluorescence. Ablation is then performed while imaging for all cells (CellTracker Red), and cut opening imaged for a further 45 s. The video concludes with a comparison of movie frames immediately after cutting vs. 45 s later.

### Supporting notes

**Note S1**. Repulsive particles pack most efficiently on flat surfaces in a hexagonal close-packed (triangular lattice) fashion where each particle has six neighbors (i.e. the coordination number *z* = 6) (**Fig. 1B**). However, this packing pattern cannot be maintained on a curved surface, since wrapping a triangular lattice onto a surface with non-zero Gaussian curvature creates strain ^96^. This strain creates an energetic cost that favors the emergence of topological defects called disclinations, in which some particles have greater or fewer than six neighbors. For example, the strain introduced into a hexagonal lattice mapped onto a spherical surface can be relieved by introducing 12 pentagons (known as topological defects of ‘charge’ *s* = +1), such as in the familiar pattern of pentagons and hexagons making up a soccer ball. For curved crystals that are sufficiently large (R/a > 5 for spheres, where *R* is sphere radius and *a* is mean particle spacing), the energy cost of isolated pentagon (*s* = +1) defects becomes too large ^18,97^. The resulting strain is therefore relieved through the spontaneous appearance of ‘excess’ dislocations as pairs, clusters, or chains of alternating charge (i.e. particles with coordination number 5-7-5-7-…-5) called ‘grain boundary scars’ each with a net charge of +1 that effectively distribute the elastic energy of the defect over a larger area ^17,18,20^.

**Note S2**. The theory of curved crystals suggests that the amount of bias toward pentagon dislocations will increase linearly with curvature. This curvature dependence of tip geometry can be summarized by plotting total disclination defect charge *Q* for individual kidney fields of view against the surface integral of their Gaussian curvature (in units of disclinations), *Ω*. The Gauss-Bonnet theorem suggests that the overall topological charge will increase linearly here to “screen” curvature-induced stress ^17,19^.

**Note S3**. Live imaging of cap mesenchyme cells in mouse kidney explants at an equivalent developmental time of ∼E14 has shown their mean speed to be approximately 3.5 × 10^-3^ μm s^-1^ (12.6 μm hr^-1^), compared to a relative divergence speed of sister UB tip domains of ∼2-5 μm hr^-1 6^. Therefore, during early branching morphogenesis, cap reorganization is sufficient to consider it as a fluid-like compartment on the timescale of tip domain duplication. Less is published on stromal cell dynamics, but for example endothelial cell induction from FOXD1+ stromal cells, recruitment, and reorganization into developing vascular plexuses at the border of tip domains is highly active during their duplication ^32^. 24-72 hr live imaging of E11.5 FOXD1+ stroma-specific reporter mouse kidney explants also suggests that cell dynamics in the stroma are sufficient to consider it to be a fluid-like compartment on the timescale of tip domain duplication (Nils Lindstrom, unpublished).

**Note S4**. Rigidity transitions have been studied in 2D vertex models (where the degrees of freedom are the vertices of sub-unit polygons) and 2D Voronoi models (where the degrees of freedom are the Voronoi centers of sub-unit polygons). The vertex and Voronoi models are distinct, but in practice the local minimum energy states of the two are very similar ^44^. Each model is based on forces imposed through the gradient of a dimensionless energy functional of the form ^42–48^:

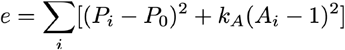

Where the average polygon area is 1, P_*i*_ is the actual perimeter of polygon *i*, P_0_ is the preferred polygon perimeter, and A_*i*_ are actual polygon areas. k_*A*_ controls the relative area and perimeter moduli. The first term models contractility and interfacial tension. For cells, this would capture cell cytoskeletal contractility and cell-cell interfacial tension, while for tip domains this would capture collective cell contractility and domain-domain interfacial tension at the cap mesenchyme-stromal interfaces (**Fig. 2B**). The second term models area elasticity. For cells, this would capture cell elasticity, while for tip domains this would capture collective cell elasticity. Rigidity transitions have been found in the 2D vertex model for k_*A*_ = 0 when the control parameter p_0_ crosses a transition point at p*_0_ = 3.87, for k_*A*_ > 0 at p*_0_ = 3.81-3.92, and in the 2D Voronoi model for k_*A*_ = 0 at p*_0_ = 3.82 (ref. ^42,46^). A strict rigidity transition does not occur in the 2D Voronoi model for k_*A*_ > 0, but it still exhibits a striking increase in e.g. elastic modulus in the vicinity of p*_0_ = 3.8 (ref. ^45^). In practice we measure the median of the observed shape index, 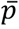, an order parameter used as a proxy for p_0_ (ref. ^42,50^). We use p* as short-hand for p*_0_ in the main text. This model family therefore identifies the shape index range of 3.8-3.92 as a threshold at which density-independent stiffening or a sharp rigidity transition may occur in tissues organized into tessellated polygonal sub-units such as cells or tip domains.

For tip domains, these energy terms are justified firstly by macroscopic behavior of embryonic kidney cortex explants, which actively contract and round up without loss of tip domain adhesion to neighbors in the hours after cutting ^16^. Secondly, elasticity of tip domains is implied by positive stiffness values returned by microindentation on a length-scale appropriate to tip domain extents in *xy* and *z* (**Fig. 3D**), and interfacial tensions are implied by rebound velocities being positive in domain-domain interface ablation experiments (**Fig. 5F**).

**Note S5**. Bayesian force inference was applied according to a recently published procedure that infers maps of tissue tension and pressure based on observed variations in edge lengths and angles at vertices of interface networks, assuming that tissues are in instantaneous mechanical equilibrium, tissue mechanics are dominated by in-plane tensions and pressures, and that interfacial tensions are positive ^62,64^. The Bayesian approach is appropriate here, since it does not require measurement of interface curvatures, making it ideal for our application to large numbers of tip domains interfaces with negligible curvatures that can be closely approximated by polygonal Voronoi cell edges ^64^, **Fig. 1C,D**. Tensions and pressures are estimated from force balances at vertices of polygon junctions, where parallel (interfacial tension) and perpendicular (hydrostatic pressure, elasticity, and contractility) force contributions are transmitted to neighboring polygons. The biological origin of these forces may be distributed throughout polygons and need not be localized at junctions themselves ^63,65^. Junctional stresses estimated through force inference are thought to be independent of other external contributions to total stress ^63^ due to e.g. out-of-plane basal interaction between tip domains and underlying cortical tissue.

**Note S6**. Other perturbations apparently unrelated to ligand secretion or spatial distribution appear to affect nephrogenesis. For example, nephrogenesis is significantly increased during embryonic kidney explant culture under surface tension at an air-media interface, rather than suspended just below the media surface or in hanging drops ^98^. Nephron formation in hemispherical bio-printed organoids is biased toward high-curvature edges ^99^, which are known to amplify mechanical strain ^87^. Nephron progenitor regulation is dependent on YAP ^90,91^, an integrator of mechanical cues affecting a range of differentiation events ^92,93^. Stromal Fat4 enhances nuclear export of YAP in nephron progenitors, tipping their perception of Wnt/β-catenin signaling toward differentiation ^72^. Finally, negative regulation of Rho GTPase, Rho-associated protein kinase (ROCK), and non-muscle myosin IIA/B isoforms (Myh9/Myh10) lead to increases in nephron formation (e.g. low dose H1152, a ROCK inhibitor) or decreases in nephron formation (e.g. high dose H1152; mesenchymal Myh9/Myh10 deletions) ^91,94,95^. Both Rho/ROCK and non-muscle myosin II are important to cell tension and perception of the mechanical microenvironment ^100,101^.

## Methods

### Animal experiments

Mouse protocols followed NIH guidelines and were approved by the Institutional Animal Care and Use Committee of the University of Pennsylvania. E14-E17 embryos were collected from timed pregnant CD-1 mice (Charles River) and stages confirmed by limb anatomy as previously described ^102^. Embryonic kidneys were dissected in chilled Dulbecco’s phosphate buffered saline (DPBS, MT21-31-CV, Corning) ^103^.

### Kidney immunofluorescence imaging

Immunofluorescence staining and imaging was performed as previously described ^16^, using protocols adapted from Combes *et al*. and O’Brien *et al*. ^104,105^. Briefly, dissected kidneys were fixed in ice cold 4% paraformaldehyde in DPBS for 20 min, washed three times for 5 min per wash in ice cold DPBS, blocked for 2 hr at room temperature in PBSTX (DPBS + 0.1% Triton X-100) containing 5% donkey serum (D9663, Sigma), incubated in primary and then secondary antibodies in blocking buffer for at least 48 hr at 4°C, alternating with 3 washes in PBSTX totaling 12-24 hours. The minimum duration of primary and secondary incubations and washes depended on the age of the kidney, as previously described ^104^.

Primary antibodies and dilutions included rabbit anti-Six2 (1:600, 11562-1-AP, Proteintech, RRID: AB_2189084), mouse anti-E-cadherin (1:200, clone 34, 610404, BD Biosciences, RRID: AB_397787), rat anti-E-cadherin (1:100, ab11512, abcam, RRID:AB_298118), mouse anti-calbindin D-28K (1:100, clone CB-955, C9849, Sigma, RRID: AB_476894), mouse anti-pan-cytokeratin (1:200, clone 11, C2931, Sigma, RRID:AB_258824), and goat anti-jagged 1 (1:150, AF599, R&D Systems, RRID: AB_2128257). Secondary antibodies (all raised in donkey) were used at 1:300 dilution and include anti-rabbit AlexaFluor 647 (A31573, ThermoFisher, RRID: AB_2536183), anti-rabbit AlexaFluor 555 (A31570, ThermoFisher, RRID: AB_2536180), anti-mouse AlexaFluor 555 (A31572, ThermoFisher, RRID: AB_162543), anti-goat AlexaFluor 488 (A11055, ThermoFisher, RRID: AB_2534102), anti-rat AlexaFluor Plus 555 (A48270, ThermoFisher, RRID: AB_2896336), anti-rabbit AlexaFluor 488 (A21206, ThermoFisher, RRID: AB_2535792), and anti-rat AlexaFluor Plus 405 (A48268, ThermoFisher, RRID: AB_2890549). In some experiments, samples were counterstained in 300 nM DAPI (4’,6-diamidino-2-phenylindole; D1306, ThermoFisher), 20 μg ml^-1^ AlexaFluor 488-labeled peanut (*Arachis hypogaea*) agglutinin lectin (PNA, L21409, Sigma), and/or 1:40 AlexaFluor 647 phalloidin (A22287, ThermoFisher) diluted in blocking buffer for 2 hours at room temperature, followed by 3 washes in PBS.

Kidneys were imaged in wells created with a 2 mm diameter biopsy punch in a ∼5 mm-thick layer of 15:1 (base:crosslinker) polydimethylsiloxane (PDMS) elastomer (Sylgard 184, 2065622, Ellsworth Adhesives) set in 35 mm coverslip-bottom dishes (FD35-100, World Precision Instruments). Imaging was performed using a Nikon Ti2-E microscope equipped with a CSU-W1 spinning disk (Yokogawa), a white light LED, laser illumination (100 mW 405, 488, and 561 nm lasers and a 75 mW 640 nm laser), a Prime 95B back-illuminated sCMOS camera (Photometrics), motorized stage, 4x/0.2 NA, 10x/0.25 NA and 20x/0.5 NA lenses (Nikon), and a stagetop environmental enclosure (OkoLabs).

### Embryonic kidney data annotation

UB tip positions were manually annotated from confocal stacks in FIJI using the *multi-point* tool. Nephrons per tip were manually annotated from confocal stacks of the SIX2 and PNA channels, summing across SIX2+ pre-tubular aggregates/early renal vesicles and later stages for which connecting tubule junctions with UB tips were visible as circular intersections in the PNA channel at tips.

Confocal immunofluorescence stacks on the SIX2 channel were manually annotated to extract kidney outlines at each *z* plane by manually tracing in FIJI on an Apple iPad. These outlines constituted a binary mask stack used for input to curvature analysis.

### Voronoi, shape index, and defect charge analysis

Delaunay triangulations and Voronoi diagrams were created from UB tip coordinates in MATLAB, filtered by polygon area and manually editing to remove edge cells. The coordination number was determined as the number of sides of each Voronoi cell, and shape index *p* was determined via *p* = *P/*√*A*, where *P* is the perimeter of a Voronoi cell and *A* is its area. Heat maps of shape index were produced by interpolation of shape index data in MATLAB using the ‘v4’ method in griddata.m. The total disclination defect charge *Q* for a particular Voronoi diagram was determined by summing the topological charges *s* over all Voronoi cells (note: *s* = 6 - *z*, where *z* is the coordination number of a cell).

### Packing model

We used physics-based modeling in the Rhino (v7, Robert McNeel & Associates) Grasshopper Kangaroo2 environment (Daniel Piker) to demonstrate repulsive sphere packing on flat surfaces and spheres in **Fig. 1B** and a kidney surface mesh in **Movie S2** (see **Kidney curvature analysis**). Quantitative principles of this software environment are detailed in ref. ^16^. We used the *SphereCollide* goal to model sphere mutual repulsion and the *OnMesh* goal to hold spheres at flat or curved interfaces, each with arbitrarily high energy potential weightings to mimic previously described features of particle packings on surfaces ^17–19,106^. To create overlay schematics, sphere coordinates were exported from Rhino and used as input to the same Voronoi and coordination number analysis performed in MATLAB.

### Kidney curvature analysis

3D mask stacks of kidney outlines were first exported from 3D viewer in FIJI as binary .stl meshes representing kidney surfaces (**Fig. S1**). These .stl files were then imported into Rhino, remeshed down to ∼1,000 polygons using quad remesh, smoothed, triangulated, draped, and split to remove the bottom face. Contour cross-sections were created at 20 μm increments along the long axis and lofted to create a smooth, regular mesh representing the outer kidney surface. Surface meshes were exported as .stl files, imported into MATLAB and represented as height maps using stlread and trisurf commands. Exported meshes were also analyzed to determine the Gaussian curvature local to each polygon via a custom workflow in Rhino Grasshopper. The area-weighted sum of polygon Gaussian curvature *K I* = *∫*_*A*_ *K dA*, over an equivalent area covered by the Voronoi diagram for that surface was then used to compute the integrated Gaussian curvature in units of disclinations, 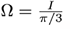.

### Spatial autocorrelation analysis

2D spatial autocorrelation heatmaps of tip regions were computed from maximum projections of confocal *z* stacks on the PNA fluorescence channel using the Wiener-Khintchine theorem in MATLAB (autocorr2d.m, Tristan Ursell).

### Brillouin microscopy

A confocal Brillouin microscope was used for acquiring mechanical images of kidneys. A detailed description of the instrumentation is published in a previous report ^107^. Briefly, the Brillouin microscope consisted of a Brillouin spectrometer and a commercial fluorescence microscope (IX83, Olympus). A continuous wave laser (∼17 mW, 660 nm, Torus, Laser Quantum) was used as light source. The laser beam was focused into a spot of 0.7 μm x 0.7 μm x 2 μm using a 40x/0.6 objective lens (Olympus). The high-resolution Brillouin spectrometer was built from a two-stage virtually imaged phased array (FSR = 15 GHz, LightMachinery). The spectrometer is shot-noise limited and has a spectral precision of 0.01 GHz, suggesting a sensitivity of 0.16% for water-based materials. The Brillouin spectra were collected using an EMCCD camera (iXon life 897, Andor). During the Brillouin experiment, kidney tip domains were scanned at room temperature with a step size of 1 μm and an acquisition speed of 50 ms per point. UB tip PNA fluorescence was also imaged on a separate channel to later register with the corresponding Brillouin images.The longitudinal modulus (*M′*) is determined by the measured Brillouin shift (*ν*_*B*_) via the relationship 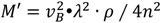, where *λ* is the wavelength of the laser, *n* and *ρ* are the refractive index and density of the sample, respectively. To calculate the longitudinal modulus, we estimated *n* = 1.40 from literature measurements of adult kidney ^108^. The density was calculated as 1.096 g/ml based on a published relationship between refractive index and density for biological materials ^54^. According to the linear log-log relationship between longitudinal modulus (*M′*) and Young’s modulus (*E*), the relative changes of both moduli are related by *ΔE*/*E* = (1/*a*) · *ΔM*/*M*, where the coefficient *a* = 0.0617 was obtained by calibration with an atomic force microscope using NIH 3T3 cells ^57^.

### Microindentation

Free-standing mouse kidneys were micro-indented immediately after explanting using established methods ^51^. Instrumentation was the same as described by Levental *et al*., except indenter arm height was controlled by a stepper motor (L4018S1204-M6, Nanotec, Munich) rather than a hydraulic micromanipulator. The indenter was fabricated from cylindrical 30 gauge (AWG) SAE 316L stainless steel wire having a diameter of 255 μm. Briefly, after calibrating the spring constant of the tensiometric sensor, each kidney was placed in a 35 mm culture dish and bulk media wicked away to leave the kidney surface semi-dry. The indenter was immediately lowered at a rate of 12.5 μm s^-1^ using the z-stage of the instrument to an indentation depth of ∼30 μm into the kidney surface while automatically recording time, force on the indenter, and indenter *z* position. Indentation was then halted and force vs. time recorded for an additional 60-120 s. Stiffness was inferred using the following relationship for indentation of a soft homogeneous material by a flat hard surface:

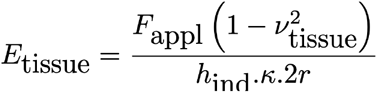

Where *F*_appl_/*h*_ind_ is the slope of force vs. indentation depth from the point of contact after accounting for sensor spring constant. *ν* is the Poisson ratio of the tissue, assumed to be 0.5. *κ* is the Hayes correction factor ^109^, accounting for finite sample thickness (*κ* was determined for each stage based on average kidney width inferred from confocal image stacks), and 2*r* is the diameter of the probe (255 μm).

Similar to previous analysis ^51^, force vs. time (*t*) traces at fixed indentation depth were normalized to maximum force and fit to the following three-parameter relaxation relationship using non-linear least-squares fitting in MATLAB (*lsqcurvefit* function):

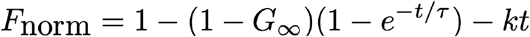

This relationship implies a Kelvin-Voigt material with an extra viscous damper in series, where G_∞_ and *τ* are an equilibrium modulus and relaxation time constant, respectively. We refer to *k* as a viscous relaxation constant in this work. For the fits presented in Fig. 3D, errors in parameter estimates were determined using the *nlparci* function in MATLAB. Stated in terms of coefficients of variation relative to the fitted parameter values (σ/value × 100) these were 0.20 ± 0.17% (mean ± S.D.) with a maximum of 0.82% for G∞, 1.2 ± 0.79% with a maximum of 3.3% for *τ*, and 0.97 ± 1.0% with a maximum of 5.8% for k.

### Force inference

Tip domain Voronoi diagrams were segmented in the *Tissue Analyzer* plugin in FIJI ^110^ and output data were passed to MATLAB for Bayesian force inference using code published by Kong *et al*. ^62,64^. We used this code to compute stress tensors and their isotropic and anisotropic (deviatoric) components for each tip domain using Batchelor’s formula ^62,64^. Since stress outputs are relative here, we normalized them to the largest median stress in a given set of comparisons.

### Laser ablation

E17 kidneys were labeled in 20 μg ml^-1^ Alexa Fluor 488-PNA and 1:1000 CellTracker Red (ThermoFisher C34552, 10 mM stock in DMSO) in Dulbecco’s minimum essential medium (DMEM, 10-013-CV, Corning) for 1 hour at 37°C, washed 3x in DMEM and placed in 2mm-diameter PDMS wells (see **Kidney immunofluorescence imaging**), this time plasma bonded to a quartz coverslip (No. 1 thickness, Ted Pella, 26014) to maximize UV transmission. Kidneys were ablated using a 355 nm UV laser (Molecular Machines & Industries, SL μCut v1.0) through a 10x objective on a Nikon Ti2 controlled by MMI software. Cut opening time-lapses were imaged from the microscope monitor using an iPhone 8 video camera and spatially calibrated using fiducial marks in the MMI software. Kidneys were fixed after ablations in ice cold 4% PFA for 15 min, washed 3x with PBS and imaged by confocal microscopy (see **Kidney immunofluorescence imaging**).

### Statistical analysis

One-way analysis of variance (ANOVA) with correction for multiple comparisons using Tukey’s honestly significant difference test was performed in MATLAB using the *anova1* and *multcompare* functions.

